# Multivariable Graphical User Interface for Simulation of Tethered Particle Motion

**DOI:** 10.1101/2022.08.31.506066

**Authors:** Khovesh A. Ramdin, Markus Hackl, Shishir P. S. Chundawat

## Abstract

The analysis of particles bound to a surface by flexible tethers can facilitate understanding of various biophysical phenomena (e.g., molecular dynamics of DNA-protein or protein-ligand binding interactions, DNA extensibility and polymer biophysics). Being able to model such systems theoretically can aid in understanding experimentally observed motions and furthermore the limitations of such models can provide insight into modeling complex systems that basic theory sometimes cannot account for. The simulation of tethered particle motion (TPM) allows for efficient analysis of complex behaviors exhibited by such systems, however this type of experiment is rarely taught in undergraduate science classes. We have developed a MATLAB simulation package intended to be used in academic contexts to concisely model and graphically represent the behavior of different tether-particle systems. We show how analysis of the simulation results can be used in biophysical research employing single molecule force spectroscopy (SMFS). Here, our simulation package is capable of modeling any given particle-tether-substrate system and allows the user to generate a parameter space with static and dynamic model components. Our simulation was successfully able to recreate generally observed experimental trends using a recently developed SMFS technique called Acoustic Force Spectroscopy (AFS). Further, the simulation was validated through consideration of the conservation of energy of the tether-bead system, trend analyses, and comparison of particle positional data from actual TPM *in silico* experiments conducted to simulate data with a parameter space similar to the AFS experimental setup. Overall, our TPM simulator and graphical user interface is suitable for use in an academic context and serves as a template for researchers to set up TPM simulations to mimic their specific SMFS experimental setup.

## Introduction

Modeling the motion of a particle attached to an extensible tether in a viscous environment is relevant to understanding fundamental biophysical phenomena as well as solving practical engineering problems. For example, the enhanced ability to observe motion of DNA scale interactions using immunofluorescence/dark field microscopy,^1^ the ability to manipulate such small-scale systems using optical/magnetic/acoustic tweezers,^2^ and the advancements in the resolution of optical imaging in the last few decades^3^ have made tethered particle analysis particularly relevant to modern-day theoretical and applied biophysics. While described in scientific literature, tethered particle motion or TPM is not typically taught in an academic context although the theory associated with this topic is crucial to understanding observations from many biophysical experiments. It is particularly necessary to study molecular-scale interactions using single-molecule experiments for comprehension of complex cellular systems which subsequently allow for improved fundamental understanding of living systems and potentially lead to the development of novel biotechnology (e.g., CRISPR enabled precision medicine).

Mathematically, tethered particle system behaviors can be approximated through the consideration of Brownian motion. Such motion is a consequence of collisions that occur between the object being tracked and the particles present in a viscous environment.^4^ In principle, fluids are composed of multiple particles that are constantly colliding. Such uncontrolled and seemingly random small-scale behaviors are better modeled stochastically since deterministic models often require unfeasible level of complexity for individual particle tracking capabilities.^5^ The idea associated with such models is to use random fluctuations to account for small scale perturbations that are observed experimentally due to diffusive effects experienced by a particle in a viscous environment.^6^

Here, we present a graphical user interface (GUI) based efficient simulation package for use by students and researchers to perform simulations of a tether-particle system with a parameter space of their choice. We have developed the underlying framework for the simulation package that builds and expands on previous models developed for educational purposes.^7^ The MATLAB-based model code was written in an easily generalizable manner, has a complete user interface and is computationally efficient so data analyses can easily be done. Simulation features like force ramps and constant force application are predefined as these are commonly encountered during SMFS experiments in real-world scenarios.^8^ Further, several corrections are included from a series of models presented in scientific literature to increase the accuracy of the TPM simulations and allow the user to understand the limitations/uses of the calculations being made. In particular, we have generated experimental data to validate our simulation predictions using a recently developed SMFS technique called acoustic force spectroscopy (AFS)^8–10^. AFS is similar to optical tweezers force spectroscopy that instead uses acoustic waves to trap micron sized particles that enable SMFS measurements for diverse tethered-particle systems. Students will be able to understand the behaviors of tethered-particle systems in general due to the easy-to-follow GUI for model presentation and exportation of several analysis plots/data from the interface to gain an appreciation for how such systems dynamically behave during live SMFS experiments.

### Experimental Setup

Our simulation experiment considers the dynamics of a bead attached to a surface using classical physics-based analysis. All the relevant parameters in this model can be altered by the user to explore alternative scenarios to aid in student learning and/or research. Further, parameters that are variable in actual SMFS experimental setups are designed to be dynamic and can be modified by the user in real-time during the simulation, mimicking an actual experiment being conducted in real-time as well. The static and dynamic parameters associated with a typical single molecule TPM system are summarized in Table 1 below:

**Table 1:** Parameters considered in the tethered particle motion simulation. Static refers to fields which are assigned before the TPM simulation starts. Dynamic refers to fields which can be modified during the simulation.

| Simulation Parameters | Parameter Notation |
| --- | --- |
| Length of Simulation (Static) | $n$ |
| Tether Persistence Length (Static) | $L_p$ |
| Tether Length (Static) | $L_o$ |
| Viscosity of Environment (Static) | $\mu$ |
| Temperature of Environment (Static) | $T$ |
| Radius of Bead (Static) | $R_B$ |
| Density of Bead Material (Static) | $\rho$ |
| Force (Dynamic) | $F = \langle F_x, F_y, F_z \rangle$ |

Based on these parameters, a complete description of the tethered bead position, applied force, intrinsic force due to particle collisions, energy, tether extension, and displacement of the bead from equilibrium are provided in the form of continually updated graphical plots. These plots are updated at a rate specified by the user in a static field prior to start of the simulation. The interface where each of these parameters are provided by the user and the key features of the simulation package are summarized in Fig. 1.

**Fig. 1:**
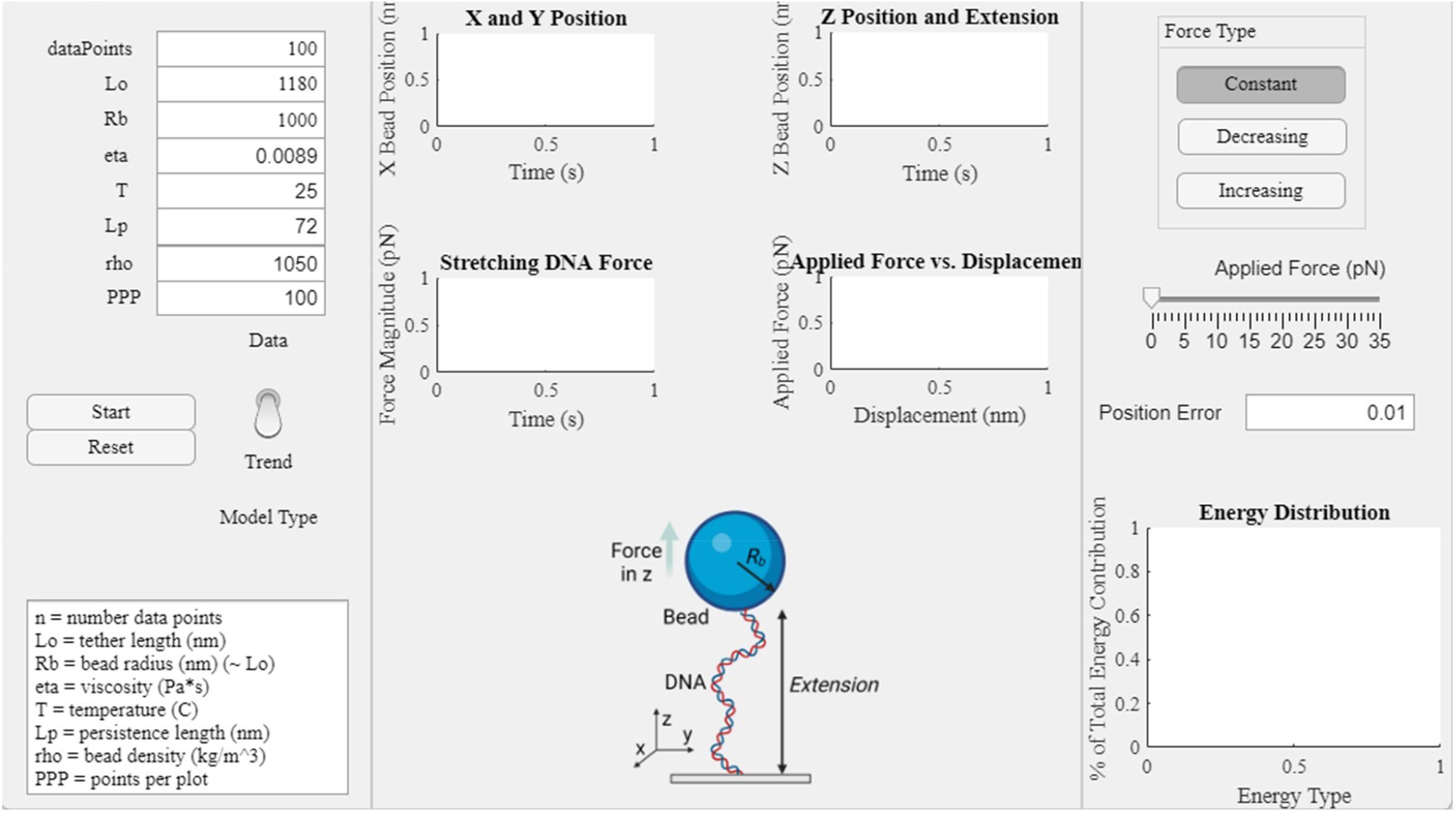
Graphic User Interface for TPM simulation at startup state.

### I. Simulation: Logic

A single MATLAB function was used that accepts user generated parameter space as well as memory terms. This function is called within a loop used in MATLAB app designer and the outputs are plotted at the user specified rate. Callback functions are utilized in the interface to synchronize the point at which the user makes a change and when that change is reflected in the base code output. The use of memory terms in the app allowed for the computations to be done with continuity as the user inputs are monitored and updated continuously within the base algorithm. That is, several simulations of the length specified by the user are run consecutively with initial conditions consistent with the end state of the prior simulation. This results in a continuous generation of data until the user specified total number of data points are reached.

### II. Simulation: Computational Framework

The notations denoted in Table 1 and Table 2 will be used in this paper to reference each variable.

**Table 2:** Summary of parameter notations used for the MATLAB code

| Parameter of Interest | Notation |
| --- | --- |
| Position (Cartesian) | x, y, z |
| Tether Extension (Displacement) | R |
| Potential Energy | PE |
| Kinetic Energy | KE |
| DNA Force | F |

The Modified Marko-Siggia worm-like chain model was considered for our model (Equation 1).^11^ Numerical root finding was used to solve for the approximate magnitude of the force for each direction.

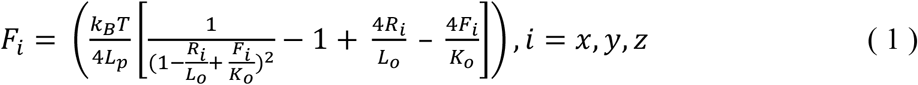

In this model, the 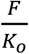 terms are a correction introduced to the classic worm-like chain model to account for the elasticity of the tether. This modification improves the experimental agreement of the worm-like chain model that only provides an order of magnitude estimate of the persistence and contour lengths.^12^ The *K*_*0*_ term is a material parameter described by Young’s modulus from classical mechanics. In this simulation, the Young’s modulus was related to the persistence length of a solid rod with a circular cross section for mathematical simplicity.^11^ The DNA diameter of d = 1.6 nm was considered in this simulation and the Young’s modulus depends on the user inputted temperature and persistence length (Equation 2). Some typical values of this parameter range from 800-1700 pN.^13,14^

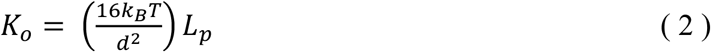

A spherical coordinate system was used to describe the particle 3D-motion in the simplest manner possible. The obtained magnitude was decomposed into x, y, and z components via projection onto a cartesian system using the following elementary trigonometric relations (Equations 3-5).

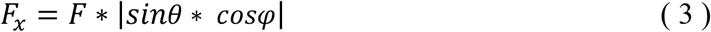

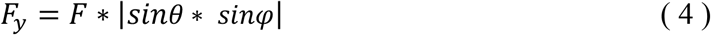

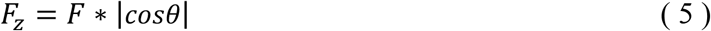

The signs of these quantities were determined directly through the consideration of the extension of the tether. If the tether was extended in a negative direction this means that the force would have to be positive to restore the system to its equilibrium position and vice versa. This described behavior is consistent with classical spring behavior described by Hooke’s Law and serves as a reasonable description for the behavior of the tether-bead system at any point in its motion due to the elasticity of the tether. All these computations were completed in a MATLAB function named ‘Marko_Siggia_Vectorized.m’ and these force computations were continuously updated in a loop from the base code. The supplementary information (SI) documentation of the simulation package provides greater detail on the functional dependencies. Users can easily grasp the purpose of each component of the simulation by reviewing the documentation of the code provided in SI and can gain a deeper understanding of the code by reading through the comments in the MATLAB files.

The computation of the *θ* and *φ* positions also come from basic trigonometric relations (Equations 6-7). The spatial orientation of the system is initially defined to be along the cartesian z direction alone and the descriptions of the angles are updates as the motion evolves over time. That is, these angular positions are derived from the consideration of the evolution of the cartesian coordinates which will later be described.

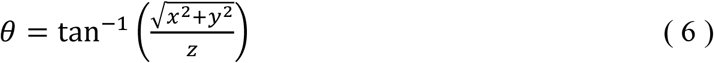

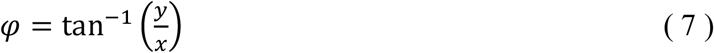

Next, the function ‘TetherForce2.m’ base code modifies the implicit force term in the z direction based on the magnitude of force applied to the system in the user interface. The first option for the user is to choose a modifiable but constant force at any point in the simulation. When the user makes a modification to the applied force using the force slider built into the app, a constant force is continually applied to the system for the duration that the user leaves the slider in the given position. This force is immediately applied with the chosen magnitude. The second option for the user is to apply a force ramp which has a slope pre-determined by the user. That is, the desired force by the end of the simulation is computed to increase in linear increments consistent with the total runtime of the simulation. The third option for the user is to apply a decaying force ramp which is computationally equivalent to the previous case except that a linear decay is considered instead of a linear increase. The projection of this magnitude onto the z direction is added to the force term from the ‘Marko_Siggia_Vectorized.m’ function to determine the net force such that *F*_*net*_ *= F*_*z*_ *+ F*_*applied*_, considering the force decomposition based on Newtons 2^nd^ Law.

The net effect experienced by the bead is intended to be consistent with Stokes Law. Namely, the bead is assumed to be perfectly spherical, all surfaces are assumed to have no imperfections, all components are assumed to be entirely homogenous, and the flow is assumed to be laminar. Laminar flow is embedded in the assumptions made since the tether-bead system is small and stable enough such that the disruptions to the environment due to the tether-bead are not as significant as the viscous forces exerted on the tether-bead system. This means that the system has a low Reynolds number which is consistent with smooth and constant fluid motion (i.e., laminar flow). A numerical validation to these assumptions is provided in the validation section of this study. Since all these conditions are approximately valid in the considered model, Stokes Law serves as a reasonable approximation to the net effect that is observed. This also means that enough information is available such that the deviations in position within a given timestep can be extrapolated from the simulation as is outlined in Equations (8) through (10) below. A correction factor is introduced to account for the edge effects since the tether-bead system is near the surface throughout the simulation. These corrections are derived based on the boundary condition that tangential flow needs to be zero at the bead surface.^15^ In accordance with classical fluid dynamics, the zero-flow condition is a consequence of the adhesive force between the fluid and surface being greater than the forces propagating motion between the fluid particles when the fluid particle is close to the surface. The x displacement (Δx) is parallel to the surface and is described in Equation 8.

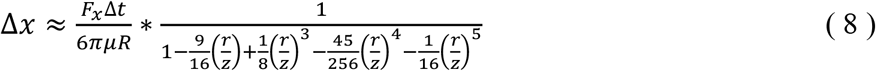

Similarly, the expressions for the y and z directions can be also obtained. The y displacement (Δy) is parallel to the surface so the correction due to surface effects remains the same. The displacement in z (Δz) is perpendicular to the plane so the correction due to surface effects is slightly different. These corrections result in the following two equations;^15^

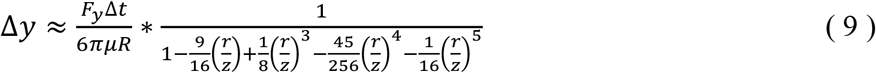

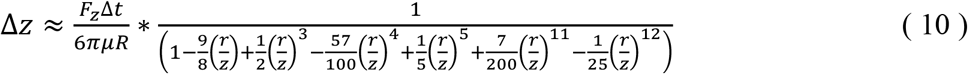

Equations (8) through (10) are used to update the x, y and z position at a given timestep. These cartesian position elements are then used in Eqs. (6) and (7) to update the spatial orientation elements from the initial state due to the viscous motion.

The rearrangement of Stokes law is only an approximation since finite timesteps are used to approximate the velocity of the bead/particle in addition to the numerical approximation of the surface effects. However, an effort was made to more accurately account for time that the particle takes to move between any two given positions through the introduction of a dynamic timestep. This dynamic timestep (Equation 11) was generated through considering the asymmetric influences that the bead experiences in the viscous environments at different extensions of the tether.^15^

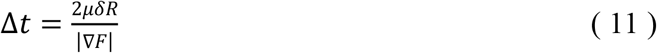

The implementation of this dynamic timestep allows for the accuracy of the position at any given time-point to be the same since the deviation is normalized using the force gradient at every datapoint^15^ This is significant because even if the users cannot extrapolate exact values at which time an observation could be made, they would be able to get a sense of the relative timescales between each observation. The modified Marko-Siggia model^11^ accounts for the extensibility of the tether. This has a direct influence on the dynamic timestep which depends on normalization using the force gradient. The explicit computations are shown in Eqs. (12) and (13) below.

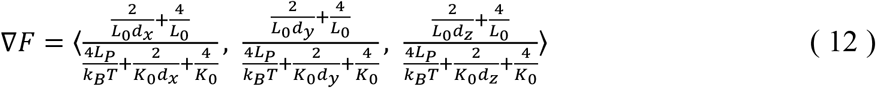

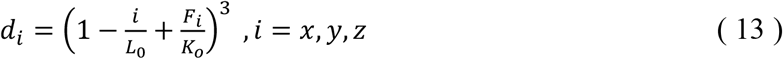

After these predictable effects are accounted for, the last remaining component necessary for accurately describing the particle position is its random motion due to diffusion. The correction factor to the position predicted from the basic force analysis was implemented using a random number generator. The random number generator was limited to having a standard deviation consistent with all the classically allowable position values obtainable by the particle and a mean value of 0 to capture the most probable trends (see equation 14). The environment is assumed to be approximately isotropic as previously mentioned, which means that the expected standard deviation is independent of direction.

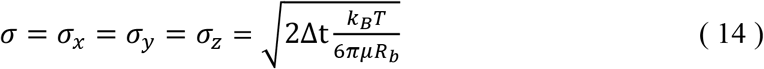

The assumptions outlined above allowed for a complete description of the position of the bead-tether system to be generated under applied force. Since the time associated with motion between any two given positions is also available, many useful computations can be done to test the validity of the code, for example the verification of the conservation of energy. The general relation *F =* −*∇U* was considered. The magnitude of the force in each direction was extrapolated between subsequent timesteps. The dynamic timestep considers the duration in which it takes for the bead to move between two points in the viscous environment explicitly so each timestep is approximated to have a force contribution independent from the next. That is, the force acting on the particle is approximated to be constant and independent for a given timestep. This means that the potential energy can be approximated using Eq. (15) below. The force terms are constant in each interval and this means that the integration only occurs over the position differential.

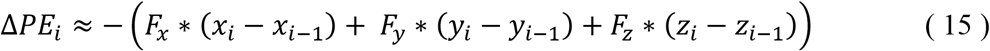

The projections of the force on the x, y, and z direction and the displacement as calculated between subsequent timesteps are considered in determining the change in potential energy. The kinetic energy was computed and rewritten using parameters relevant to the constructed system in Eq. (16) below.

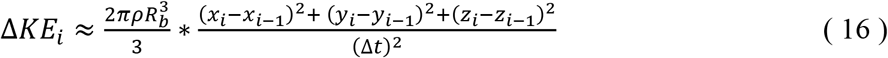

The *ρ*term in the kinetic energy computation is the density of the bead considered in the system, the displacement in each direction is determined at subsequent timesteps iterated by variable i and the length of the timestep is denoted *Δ t*.

All the arguments made above are consistent with a probabilistic consideration of the behavior of the tether-particle system. This means that by nature, assumptions of thermal equilibrium are made as can be seen from the applications of the equipartition theorem. These assumptions are not appropriate when a force is instantly applied to the system. The force ramp feature utilized allows for a steady buildup of the force which does not perturb the system by a great extent at any given instant. However, for general applications of large magnitudes of force, this model can break apart. As such, a separate model was implemented which uses physical constraints to ensure that the system remains stable as expected in reality.

First, in the cases where a force is applied to the system, an approximation such that *F*_*net*_ *≈F*_*applied*_. When a force is applied, the tether will extend, and the tension will increase resulting in limited fluctuations. In accordance with the modified Marko-Siggia model,^11^ these fluctuations will occur about an equilibrium value which is associated with the applied force. Extracting this equilibrium value allows for a description of the bead position to be made independent of the timescale associated with the instability. Generating a distribution of permissible values about this equilibrium position allows for a complete description of the bead position. The values attainable by the bead are explicitly constrained by the tether length and the small probability of the bead reaching a value significantly different from the equilibrium value. These constraints are well described by a normal distribution with 0 mean fluctuations about the equilibrium position. The standard deviation was determined as shown in Eq. (17) below where *α* is an arbitrary parameter meant to describe the resistivity of the environment. This parameter is not strictly defined and can be modified to best fit the data collected. In accordance with the assumption of isotropy, *α = 3* was assigned as the base setting.

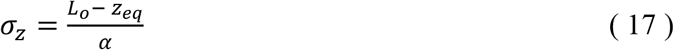

Since the z spatial harmonic behavior is described, the planar region of interest can easily be extrapolated. The system is defined such that magnitude of the position vector corresponds to the tether extension. Since a reasonable approximation to the z position is obtained, the acceptable x and y positions must be approximately consistent with the constraint in Eq. (18) since a force regime in which unwinding of the double stranded DNA occurs (∼ 65 pN) is not considered here.

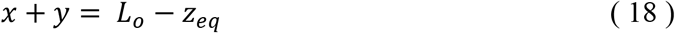

The distribution of the x and y positions are not expected to have significant bias since an isotropic environment is considered. As such, the weight of the permissible positions will be approximated to be equivalent.

Using these two conditions, a constraint for the x and y position can be obtained. Since the viscous effects also have a contribution to the planar variations, a distribution was generated under the assumption of thermodynamic equilibrium in Eq. (14). The instantaneous application of force in the z direction is accounted for without the consideration of thermodynamic equilibrium. Essentially, this means that the system is forced into a harmonic state which can again be described by conditions using thermodynamic equilibrium. The implementation of the previously described method is a boundary condition which restabilizes the environment. This means that the assumption of thermodynamic equilibrium is valid after the system is constrained with this method. This equilibrium is artificial in the sense that constraining the system requires higher force fluctuation magnitudes than is experimentally observed. A spatial resolution of 10% was used to limit the simulation to values similar to experimental fluctuations. This is an over approximation of experimentally observed fluctuations, but the simulation only provides an order of magnitude estimate. As a consequence, it is unfeasible to obtain results with greater precision at present. Eqs. (19) and (20) below describe the x and y positions of the bead within this framework. The computations done in the simulation begin by considering the origin of the system at the frame of the bead. All the terms were rescaled such that the result is consistent with these relations.

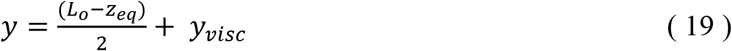

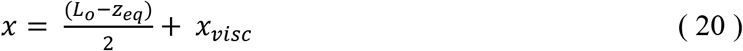

The assumptions made when generating this simulation are consistent with the assumptions made in a typical Markov process.^16^ The distributions from which the Brownian fluctuations are determined are normal with a mean of 0 and standard deviation consistent with Eq. (14). The normal distribution is a continuous probability function meaning that the area bounded by the curve is 1. Next, all of the Brownian fluctuations are extrapolated from probability distributions governed by the same rules, so all of the terms are determined using the same probability constraints. Each distribution generated at each given timestep is time independent. The dynamic timestep provides an intrinsic constraint on how far a particle could be displaced due to a collision with the molecules in the viscous environment. This means that at any given position, the span of reasonable values classically obtainable by the particles is predefined for a given timestep. Last, each state attained by the particle is assumed to be independent of every subsequent state obtained by the particle. This is consistent with the constraints associated with classical Brownian motion.^17^ Discrepancies are present due to the lack of consideration of the time-lagged covariances^18^ that were neglected since the effect is irrelevant to the level of precision considered in this simulation. There is indication that the modified time-step based on the extensibility gives a reasonable approximation to the timestep since the simulation does not attain physically unreasonable values or have a time which approaches infinitesimal scales when a force is applied using a parameter space consistent with all of the assumptions described here.

### III. Simulation: Implementation and Theoretical Validation

All the data is generated using two primary tiers of code. The base code, TetherForce2, runs the simulation based on the equations previously described using for loops. Next, a loop in the MATLAB App Designer was used to continually update the arrays containing the parameter space generated. The results generated in the base code based on the updated parameter space are immediately assigned to relevant GUI axes to plot the results as the simulation continues running in real-time. This second tier of code is the most inefficient component of this simulation since it must check for user input at every datapoint, update the parameters the base function calls for at every datapoint, and plot the data at a rate specified by the user. A more comprehensive discussion of the simulation efficiency is provided in the SI documentation. In case the code is being used solely for data generation (and not GUI based results visualization in real-time), the user can set the plot rate equal to the total number of datapoints and this will result in significantly improved runtime. While slightly more computationally intensive, it was found that continual application of a force did not significantly affect the runtime of the simulation.

Aside from the actual implementation, the simulation gives a reasonable approximation to the physical behaviors associated with a typical tethered particle-bead system. That is, the tether particle-bead system will display a wide range of fluctuation in every direction provided that an external force is not applied to the system. This is a consequence of the fact that the system will be consistently colliding with the particles of the chosen environment and will have small scale motion due to these collisions. Due to the lack of tension on the tether in the absence of an applied force, the planar variation will be more prominent since the tether-bead system will not have any rigidity as shown in the upper panel of Fig. 2 below.

**Fig. 2:**
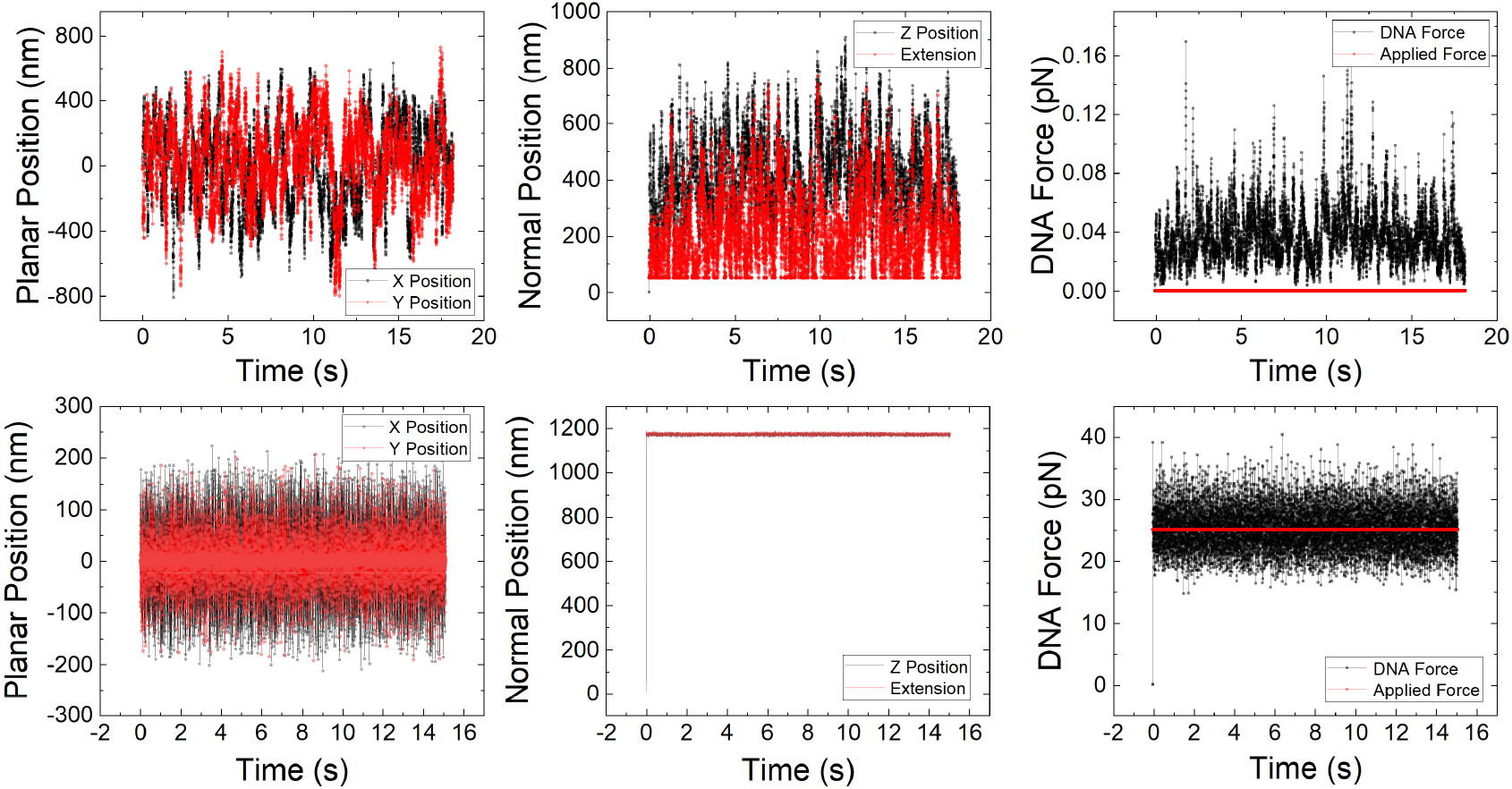
Simulation model predictions for TPM with or without applied force on system. Upper Left Panel: Planar position (XY) vs. Time in absence of applied force. Upper Middle Panel: Normal position (Z) vs. Time in absence of applied force. Upper Right Panel: DNA Force vs. Time in absence of applied force. Lower Left Panel: Planar position (XY) vs. Time in presence of 25 pN Applied Force. Lower Middle Panel: Normal position (Z) vs. Time in presence of 25 pN Applied Force. Lower Right Panel: DNA Force vs. Time in presence of 25 pN Applied Force. All panels were generated using the trend mode of the simulation GUI.

In the presence of an applied force, it is expected that the particle will asymptotically approach maximum extension over a given time and the x-y motion will be confined to smaller scales due to the tension experienced by the tether. The lower panel of Fig. 2 was generated using the trend mode of the simulation. The details of this mode will be discussed in more detail below. The parameter space used to generate the results plots are outlined in Table 3.

**Table 3:** Sample parameter space used to generate Fig.2 result plots.

| Parameter (unit) | Value |
| --- | --- |
| n (-) | 10000 |
| $L_o$ (nm) | 1180 |
| $L_p$ (nm) | 72 |
| $\mu$ (Pa*s) | 0.0089 |
| T (°C) | 25 |
| $R_B$ (nm) | 50 |
| Plot Rate | 10000 |

In the absence of an applied force, the planar position of the particle is unrestricted and varies with a span of approximately ±800 nm. In the presence of an applied force, the span in which the planar position varies is limited to approximately ±200 nm. This behavior becomes more evident as the magnitude of the force increases over time since the tether will experience increasing tension. In presence of an applied force, the system asymptotically fluctuates near maximum extension. All these behaviors are consistent with theoretical expectations.

A small sample size of 10,000 datapoints is considered to capture the local trends for Fig. 2 but more valuable information of the evolution of the TPM system can be obtained via utilizing the simulation. Features like the dynamic timestep and manual variation of the force cannot be appreciated by considering a fixed plot of the behaviors associated with the generated system.

The final features implemented into this simulation are analysis plots. The relative contributions of the potential and kinetic energy of the tether bead system are useful even though a complete description of energy conservation cannot be provided. Although the total energy of the system is conserved, the energy imparted onto the environment is not trivially obtainable, so a complete conservation analysis is not considered. Rather, a consideration of the distribution of the kinetic and potential energy gives insight into the validity of the chosen parameter space. This model is generated under the assumption that the viscous forces dominate the forces exerted due to the particle on the environment. That means that this model serves as a reasonable approximation if the potential energy of the tether-bead system dominates the kinetic energy. For the system considered in this set of validation runs, this expectation is verified in the left panel of Fig. 3 below. The right panel of Fig. 3 presents a case where the simulation results are not valid due to the low viscosity. Users can use these plots to ascertain if the simulation provides a reasonable approximation for their parameter space.

**Fig. 3:**
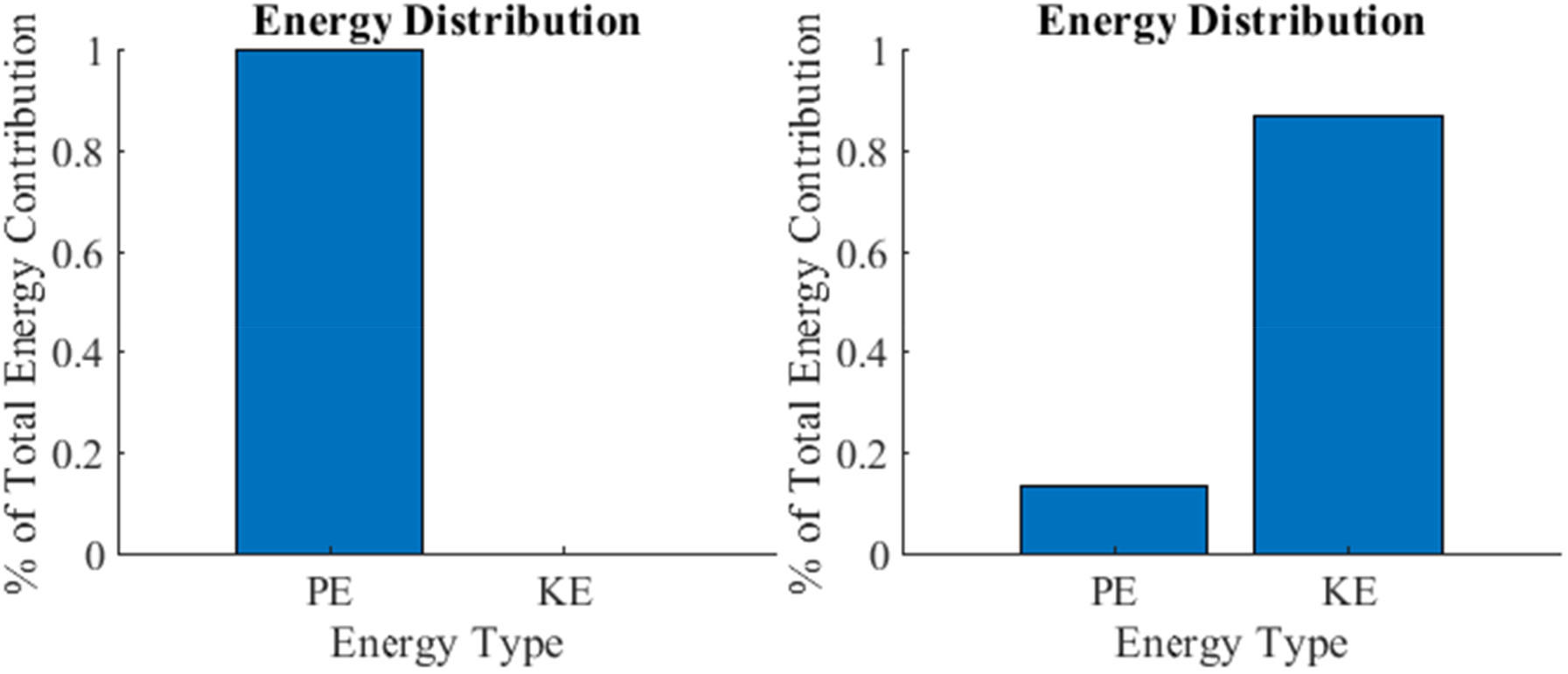
Potential and kinetic energy of the tether bead system based on the TPM simulations. Left Panel: Energy distribution for viscosity = 8.9e-4 Pa*s. Right Panel: Energy distribution for viscosity = 2e-6 Pa*s. All other simulation parameters remained the same.

Another useful analysis which validates the results of the simulation is the consideration of the applied and intrinsic DNA forces as a function of the displacement. In this model, the maximum extension should not greatly exceed the combined length of the bead and tether at any timepoint since the force application is limited to a regime in which dsDNA helix unwinding is not relevant as previously discussed. Fig. 4 below confirms this physical restriction both in the presence and absence of an applied force.

**Fig. 4:**
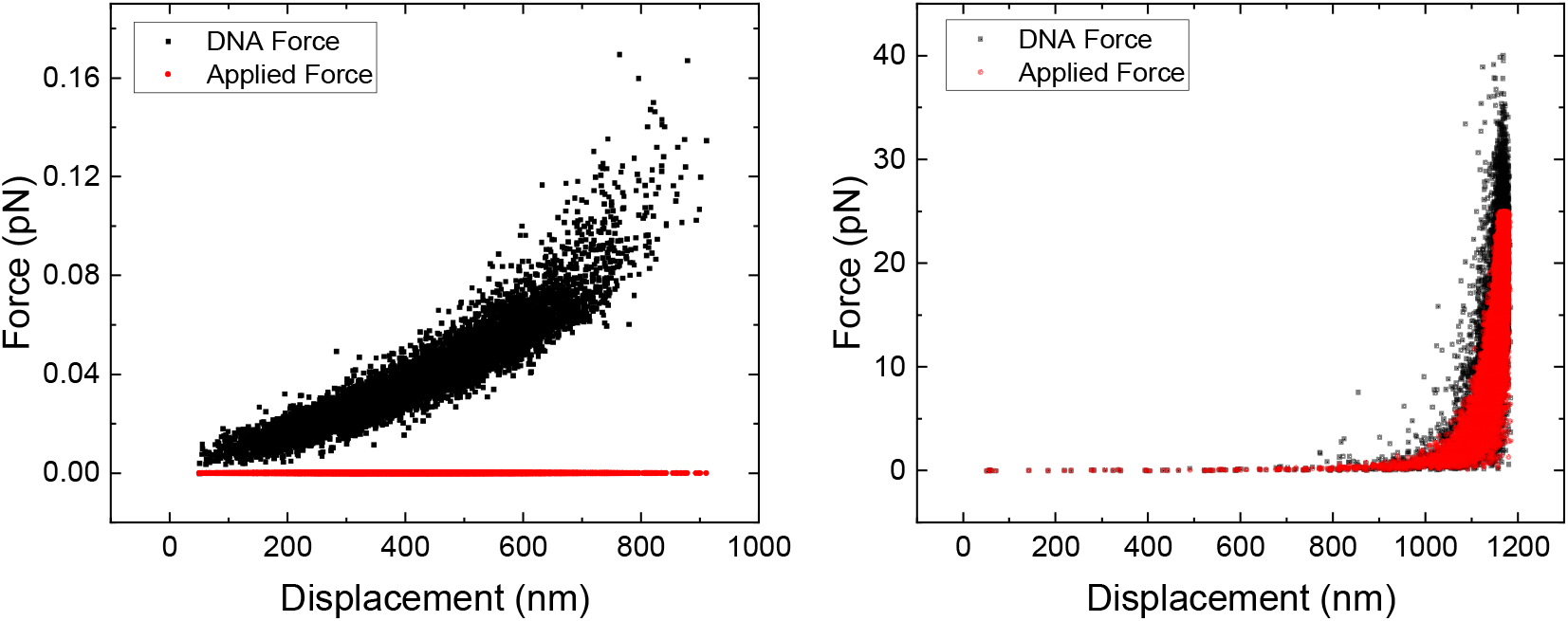
Force Displacement Curves in Absence of Force (left) and in Presence of 25 pN Force Ramp (Right).

### IV. Simulation: Model Validation Via Acoustic Force Spectroscopy Experiments

For the comparison of the simulation results with SMFS experimental data, a TPM experiment was carried out in our laboratory using an Acoustic Force Spectroscopy (AFS) instrument.^9,19^ A more detailed description of the AFS experimental setup is found in the SI document and is also described in our recent publication^10^. Briefly, the surface of the AFS chip is incubated with anti-digoxigenin fab fragments for 20 minutes, followed by a surface passivation with bovine serum albumin (BSA) protein, casein protein, and Pluronic F-127 nonionic surfactant in 10mM phosphate buffered saline (PBS) solution for 30 minutes. Next, DNA functionalized on opposite ends with biotin and digoxigenin is mixed with streptavidin-functionalized polystyrene beads for 30 minutes, washed twice in PBS containing BSA, casein, and Pluronic F-127 and incubated in the AFS imaging chip for 15 to 30 minutes. Finally, non-bound beads are flushed out at a flow rate of 2μl/min and the remaining beads are tracked in 3D. Analysis of bead traces was performed with the software provided by LUMICKS, and a free academic version can be found in the original publication of the AFS.^19^

Next, some results from actual TPM experiments conducted in this project using the AFS are presented to verify that the simulation provides an order of magnitude estimate to expected x/y root-mean-square position values.

As can be observed in Fig. 5, the simulated RMS position values of the bead-tether systems are of the same order of magnitude and follow the same trend as the AFS experimental values. The RMS value^19^ was determined using Eq. (21).

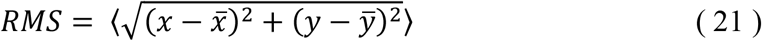

**Fig. 5:**
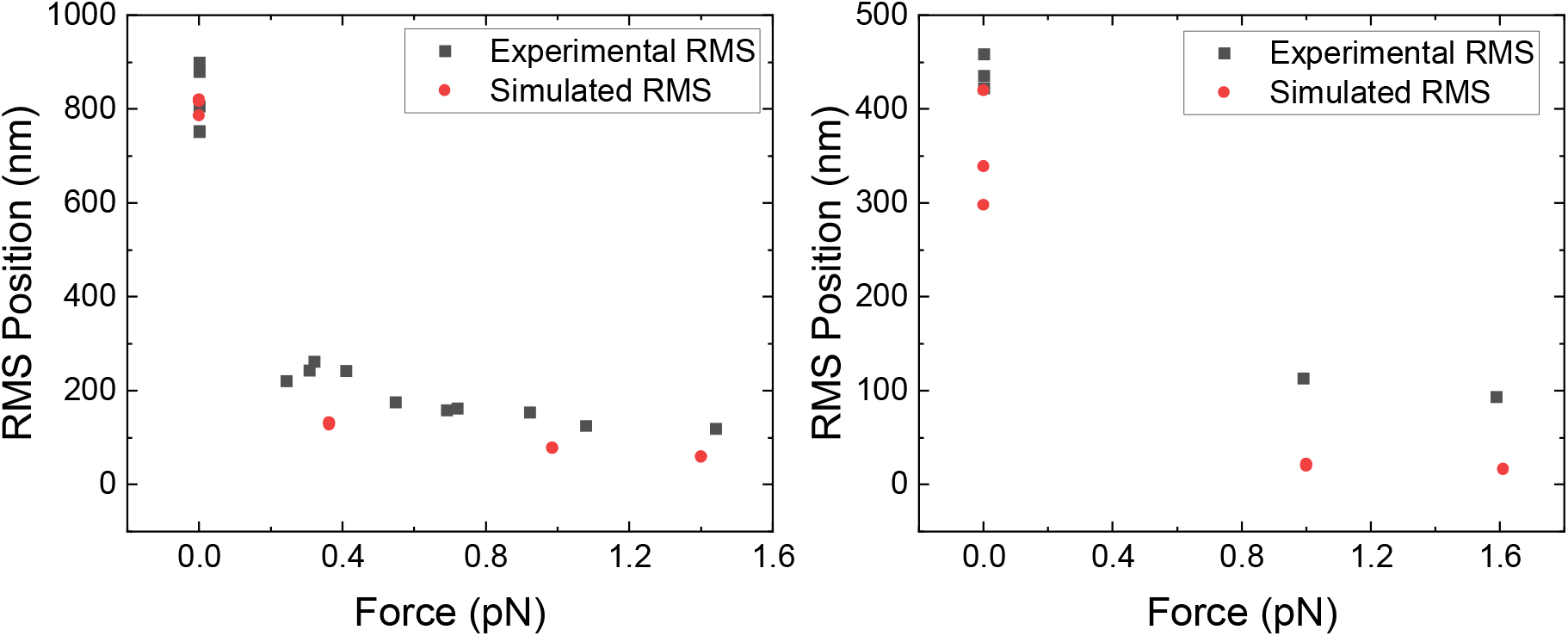
Simulation model agrees well with TPM observed in Acoustic Force Spectroscopy experiments. Left Panel: Experimental and simulated average x/y-RMS position values with application of constant force for 1800 nm DNA strands attached to 3110 nm polystyrene bead in water at room temperature. Right Panel: Experimental and simulated average XY-RMS position values with application of constant force for a 500 nm DNA strand attached to a 3110 nm polystyrene bead in water at room temperature.

As the magnitude of applied force increases, the equilibrium RMS position the system takes also increases. The greater variation in AFS experimental RMS values is due to experimental limitations such as multiple tether binding events, uncertainty in the true length of the tether, uncertainty in the constancy of force application as well as measurement uncertainty. It was particularly difficult to obtain beads bound by the tether for the 500 nm tether due to issues with the sample preparation. Even so, with the limited AFS experimental dataset, the TPM shows trends on the same order of magnitude as the simulation. In all these simulations, the bead radius was set at 3110 nm. The bead radius does not influence the harmonic behavior and only affects the scaling of the time from the earlier described dynamic timestep. Since this model was constrained by the tether length, bead radius values like the tether length are better at attaining approximations consistent with experiment since the scaling in time is more trivially obtained.

## Discussion

In its current setup, our simulation package provides an efficient means of generating an estimate for how a tethered particle-bead system behaves in a viscous fluid environment. The introduction of the extensibility of the tether into the worm-like chain model accounts for the elasticity of DNA when forces are applied.^13^ This means that while the parameters of the simulation need to be tuned by the user to model the system that they are studying, the general behaviors are within a reasonable number of limits that can be extrapolated for any given system. However, there are some computational limitations associated with this model. That is, a basic reduction reveals that the force model has multiple solutions. As the simulation runs, the numerical solver tends to choose a solution which can best numerically minimize the equation. While numerically reasonable, the alternate solution is not physically meaningful. All the descriptions made in the initial model are developed under the assumption of thermal equilibrium. When a force of sufficient magnitude is applied, the timestep calculation which depends inversely on the force gradient becomes infinitesimally small. This results in the computations of the position terms becoming unreasonable as well since position terms depends on the timestep. Consequently, a new model was implemented to preserve the validity of the assumptions of thermodynamic stability throughout the simulation. Specifically, two models were implemented into our simulation interface: trend analysis and data collection only.

First, the trend model captures the overall behaviors exhibited by the system. There are some numerical inconsistencies which exist in the model. In transitioning between a state of stability and artificial stability, the planar position will depict a slight increase in magnitude of fluctuations. This is consistent with expectation since the constraints inherently require the system to achieve a length consistent with the tether length. This means that the algorithm utilized maximization to generate a reasonable span of values which could be attained by the particle. The values attainable by the particle in the probabilistic approach tended to be closer to the equilibrium state but the span was not so strictly constrained. These methods represent different forms of numerical approximations and these discrepancies become evident as the simulation runs. The overall trends are still captured by the simulation and the discrepancies provide some insight into the nature of the system. That is, the physical constraints provide an upper limit to the acceptable planar position values attained by the system whereas the probabilistic approach represents the average expected fluctuations. For the purposes of strict data collection, the data model has been implemented such that the physically constrained model is used for the entire data collection period, eliminating the transition point.

A more precise model would require consideration of the bending of DNA beyond the persistence length considered in the worm-like chain model. The bending of the molecular structure of DNA on smaller scales than the persistence length is an experimentally known fact and the worm-like chain model does not account for this.^20^ Theoretically, this phenomenon can be accounted for through the consideration of the elastic collisions between the molecular bond sites and photons which result in small scale bending. This effect is known as Raman Scattering and it can be modeled through the consideration of MD and QM/MM modeling techniques.^21^ While possible, such a simulation is extremely computationally intensive, and the only correction provided to the worm-like chain model is at small scales. DNA on longer length scales have behavior well approximated by the current semi-flexible polymer description of the modified worm-like chain model since the average of the individual base-pair contributions are well approximated by a single persistence length. Correcting for smaller characteristic length scales by identifying each Raman peak does not add value to this current simulation model. The purpose of this simulation is to give students order of magnitude estimates of the expected experimental behavior of the system. While such an effect is also detectable, it is not experimentally feasible to observe DNA on such small scales without it tangling, meaning that the trends that would be observed by such considerations are still largely theoretical.^21^ For a general user trying to understand how a tether-particle behaves, the consideration of DNA as a semi-flexible polymer with a single persistence length is sufficient in most cases.

For general cases, we have shown there is reasonable agreement between the predictions of the TPM simulation model and AFS experimental results. Studies have been conducted which attribute anisotropies in single molecule TPM measurements to specific protein functionality.^22^ While such effects are experimentally observable, there is no general way of accounting for the binding dynamics of all protein and tether combinations. This simulation is written in a manner where specific systems can easily be considered. All the parameters associated with the system construction are modifiable in the current user interface. Further, the implicit construction of the bead and tether are done through experimentally modifiable quantities such as material density, stress tolerance, etc. This means an extension can easily be made to transform the bead considered in the current model into a specific protein, perhaps through the implementation of a bead subfunction which accounts for the specific characteristics relevant to the protein-ligand system of interest. Using such a subfunction in combination with a subfunction that considers DNA-Protein complex formation would allow for a completely general description of the protein-DNA binding behavior as well. As previously mentioned, the Marko_Siggia_Vectorized subfunction also provides a way for the user to easily modify the expected DNA force fluctuations by implementing corrections that more adequately account for specific systems of study. Overall, this simulation is a template that can be easily generalized to be made relevant to any specific particular area of instruction or research.

From an educational context, the simulation presented here is diverse enough to promote student learning. The code is written such that the student can follow through the logic and go through the derivations done to solve for all the outputs from first principles. All the assumptions made are outlined in the comments of the MATLAB code. While a trial study has not been conducted using this simulation in an actual classroom setting, several studies have been conducted to determine the effects of using computational sciences and simulations in a classroom setting to promote student STEM learning. Allowing students to work through simulations on their own rather than offering step-by-step guidance is often observed to result in better learning outcomes.^23^ In some similar studies, the use of simulations were found to promote knowledge integration processes, which implies that students were able to form a deeper level of understanding of the material due to their exposure to the material.^24^ The results for these types of studies tend to be diverse due to the extensive number of confounding variables present in such trials. However, these studies tend to compare the effectiveness of simulation-based vs. classical instruction-based learning and it is widely found that both provide similar results for evaluating pedagogical effectiveness. This simulation package with an easy-to-follow GUI was created with the intention of providing students and instructors the opportunity to quickly review a highly relevant topic in modern physics & engineering in a very short amount of time that would otherwise not be covered.

The efficiency of this simulation was validated through time averaging a repetition of 5 trials for all the plot rate-datapoint combinations previously considered. The consistency of the simulation in the trials has verified that a few minutes are sufficiently long enough to extrapolate all the noticeable trends associated with a tether particle system suggesting that such simulations could reasonably be used within a typical instructional lecture period. Further, the students could be tasked with using the code to do more detailed analysis such as model fitting for the data since the code outputs all the generated data as text files or studying the effect of various system parameters on experimental observations. Questions could also be asked about the logic used to develop the model as the manipulations made are clearly defined. Elementary knowledge of Physics and Trigonometry is all that is necessary to follow the logic for early undergraduate students even if they cannot understand the finer details. Upper-level undergraduate students should be able to follow the logic and derive every relation considered using the given models. The use of force-extension curves, RMS position analysis, etc. for data analysis are highly valuable tools for students to gain an understanding of tethered-particle systems.

## Conclusions

This work describes a simulation for tethered particle motion which comes with a customizable user interface. By utilization of the modified worm-like chain model with suitable corrections due to geometrical constraints, a simulation capable of predicting trends consistent with experimentally obtained data of tethered particle motion was shown here. Using the MATLAB app designer, a user interface was created which allows users to interactively modify parameters in the simulation as they would be able to do if they were conducting an actual SMFS or TPM experiment. Trials were setup to validate our simulation through consideration of behaviors in limits, verification of assumptions made, and comparison to actual experimental data. All these tests indicated that the simulation provides a reasonable classical description of tethered particle motion.

Some fundamental limitations of the simulation are associated with the bead size, the instability in the probabilistic approach and the consideration of a fixed persistence length. The bead size being constrained is not problematic since the harmonic behavior is not affected by it generally. The simulation may have some errors if the bead size is unreasonably large or small relative to the tether length, but the user can modify their system to account for this deficiency without any loss of accuracy. The choice of bead values similar to the tether length, as in the default value placed in the simulation, will allow for any experiment to be replicated. The instability in the probabilistic approach was accounted for using the constraint model. Lastly, implementing a model better able to describe the bending properties of DNA would require much more computationally intensive simulations which are intrinsically inefficient and therefore contradicts the goal of this simulation. Tests of efficiency were conducted to verify that the simulation can be used within an educational context to quickly but effectively allow students to understand this important topic and generate data within the time frame of a typical academic lecture. Having a sense of the dynamics of DNA-scale or similar polymer-tether systems will allow students to gain an intuitive understanding and insight into what can be expected from single molecule experiments using advanced techniques like optical tweezers or acoustic force spectroscopy. As advanced imaging techniques based experimental tools gain more traction both in the real-world (e.g., point-of-care diagnostics) and academic world (e.g., single-molecule imaging of cellular biophysical phenomena), it becomes imperative to expose students/researchers early on to such techniques in a classroom setting with an appropriate simulation toolkit.

## Supporting information

MATLAB files

## Acknowledgments

Prof. Shishir P.S. Chundawat acknowledges support from the Rutgers Aresty Research Center and the National Science Foundation (NSF CBET Award 1846797), Khovesh Ramdin acknowledges Rujuta Mokal for reviewing logic and generally providing support in the development of the code and writing of the paper. The DNA-bead sketch in the GUI was generated using Biorender.com.

## References

1 S. Brinkers, H.R.C. Dietrich, F.H. De Groote, I.T. Young, and B. Rieger, J. Chem. Phys. 130, (2009).

2 I. Heller, T.P. Hoekstra, G.A. King, E.J.G. Peterman, and G.J.L. Wuite, Chem. Rev. 114, 3087 (2014).

3 R. Jungmann, M. Scheible, and F.C. Simmel, Wiley Interdiscip. Rev. Nanomedicine Nanobiotechnology 4, 66 (2012).

4 K.A. Floyd, A.R. Eberly, and M. Hadjifrangiskou, Biofilms Implant. Med. Devices Infect. Control 47 (2017).

5 S. Chandrasekhar, Dialectica 3, 114 (1949).

6 S. Kumar, C. Manzo, C. Zurla, S. Ucuncuoglu, L. Finzi, and D. Dunlap, Biophys. J. 106, 399 (2014).

7 J.F. Beausang, C. Zurla, L. Finzi, L. Sullivan, and P.C. Nelson, Am. J. Phys. 75, 520 (2007).

8 G. Sitters, N. Laurens, E.J. De Rijk, H. Kress, E.J.G. Peterman, and G.J.L. Wuite, Biophys. J. 110, 44 (2016).

9 D. Kamsma, R. Creyghton, G. Sitters, G.J.L. Wuite, and E.J.G. Peterman, Methods 105, 26 (2016).

10 M. Hackl, E. V. Contrada, J.E. Ash, A. Kulkarni, J. Yoon, H.-Y. Cho, K.-B. Lee, J.M. Yarbrough, and S.P.S. Chundawat, BioRxiv 2021.09.20.461102 (2021).

11 M.D. Wang, H. Yin, R. Landick, J. Gelles, and S.M. Block, Biophys. J. 72, 1335 (1997).

12 C. Epstein and A.. Mann, Nat. Commun. 14, (2012).

13 S.B. Smith, Y. Cui, C. Bustamante, and S.B. Smith, 271, 795 (1996).

14 O.D. Broekmans, G.A. King, G.J. Stephens, and G.J.L. Wuite, 2, (2014).

15 E.W.A. Visser, Biosensing Based on Tethered Particle Motion (2017).

16 G.T.D. Kampen N.G. Van, Ibe C. Oliver, Pinsky A. Mark, Karlin Samuel, Science (80-.). 1 (2007).

17 S. Chandrasekhar, Rev. Mod. Phys. 15, 1 (1943).

18 T. DelSole, J. Atmos. Sci. 57, 2158 (2000).

19 G. Sitters, D. Kamsma, G. Thalhammer, M. Ritsch-Marte, E.J.G. Peterman, and G.J.L. Wuite, Nat. Methods 12, 47 (2014).

20 P. Wiggins, T. Van Der Heijden, F. Moreno-Herrero, A. Spakowitz, R. Phillips, J. Widom, C. Dekker, and P.C. Nelson, Nat. Nanotechnol. 1, 137 (2006).

21 S. Rao, S. Raj, B. Cossins, M. Marro, V. Guallar, and D. Petrov, Biophys. J. 104, 156 (2013).

22 Y. Ishii, Y. Taniguchi, M. Iwaki, and T. Yanagida, BioSystems 93, 34 (2008).

23 K.E. Chang, Y.L. Chen, H.Y. Lin, and Y.T. Sung, Comput. Educ. 51, 1486 (2008).

24 R. Taub, M. Armoni, E. Bagno, and M. Ben-Ari, Comput. Educ. 87, 10 (2015).

