## Supplementary Information for "Multivariable Graphical User Interface for Simulation of Tethered Particle Motion"

**Table of Contents for SI Document:**

1. Documentation for TPM simulation model package

2. Supplementary Information Results

3. Supplementary Information Methods

Supplementary Information References

### 1. Documentation for TPM simulation model package

#### Definitions

**Batch Processing**: Each simulation length is determined by the parameter n, which represents the total number of data points. The data points are generated in batches continuously and appended into a single output array based on the user specified points per plot until the $n^{th}$ data point is reached.

**Points Per Plot**: The number of data points being simulated per batch are referred to as points per plot. Each batch is plotted when the data points for that given batch is done being simulated by the base function TetherForce2. This process is continued until the user specified total number of data points are reached.

**User Specification:** Using the graphic user interface provided, the user can input parameters of their choice to be simulated. Except for the Force Slider, all of the fields must be specified prior to the start of the simulation.

**Memory Terms**: The overall simulation is run using the user specified parameters. The batches in which each of the simulations occur have initial conditions based on the end state of the prior batch simulated to ensure continuity. For the first batch produced, the initial conditions are set such that the planar position is at 0 and the normal position is at the bead radius, Rb. All of the force contributions and time are also initialized to 0.

**Planar Position:** This is the plane containing x-y values from a cartesian reference frame.

**Normal Position:** This is the plane perpendicular to the x-y plane (z direction) from a cartesian frame.

**Graphic User Interface:** The terminal the user will interact with to produce their desired simulation. This includes fields that allow for user specification in addition to different graphic outputs which summarize the simulation results.

#### Base Function

**TetherForce2**: This function accepts user specified parameters from the graphic user interface and the memory terms as input parameters. Each simulation is run for the user specified batch size. This function outputs arrays containing the planar and normal positions for each data point, the time elapsed since the start of the simulation for every given data point, the x, y and z components of the forces acting on the bead due to collisions between the particle and environment and the extension, theta and phi positions of the bead from a traditional spherical coordinate system.

#### Subfunctions

These are used to increase the readability of the code. They require inputs from the base function, tend to perform some form of extended computation and output results needed in the base code for the simulation to continue running.

**getNextTimeStepmod:** This subfunction accepts all parameters necessary for the computation of the time-step. Some of these include user-specified parameters and others include parameters obtained prior within the same simulation. For greater discussion of the logic, specific parameters used and manner of computation, reference the manuscript associated with this work.

**Marko_Siggia_Vectorized:** This subfunction utilizes a numeric solver to obtain force values based on the modified Marko-Siggia worm like chain model (1) for each direction in a cartesian system. The input parameters for this subfunction are described explicitly in the manuscript. An addition made to this subfunction is the projection of the force terms based on the theta and phi parameters. These can be added to the simulation as dynamic parameters but were neglected due to time constraints for this project. The details of the accepted parameters are presented in the manuscript associated with this work.

**getDragCoefXY:** This subfunction outputs the planar drag coefficient based on boundary conditions constraining the system. A more through discussion can be found in the manuscript.

**getDragCoefZ:** This subfunction outputs the normal drag coefficient based on boundary conditions constraining the system. A more through discussion can be found in the manuscript.

#### Graphic User Interface:

**Callback Functions:** These callbacks have user defined functions embedded in them which perform a task based on given input. For instance, the user interface has a start button which initiates the simulation based on the user inputted parameters and a reset button which clears the generated event

**Features:** The simulation includes numeric edit field for the static parameters, a toggle switch for the model type, two buttons for the start and reset functions, a set of 3 linked buttons of which only one can be chosen at a time, a slider for the dynamic editing of the force during runtime and plots of relevant parameters. A more in-depth description of how each function was implemented is presented as comments in the code and throughout the manuscript.

Overall, the flow chart in Fig. S1 below summarizes how these components interact with each other.

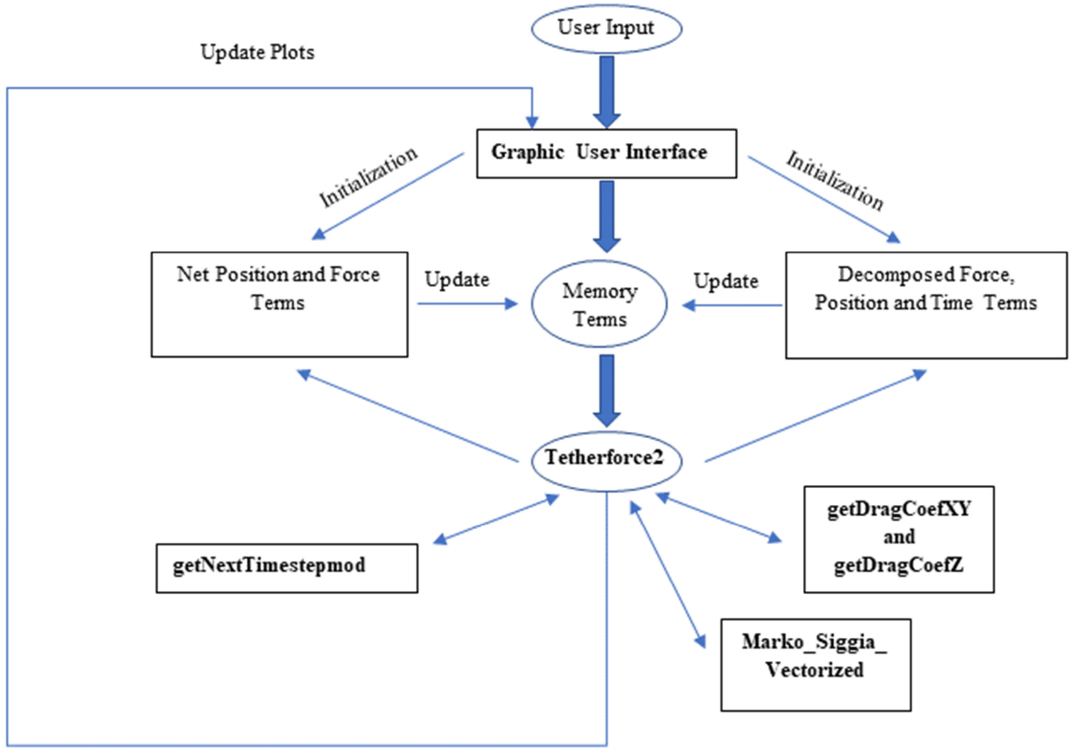


*Fig. S1: Flow chart with functional dependencies and model update logic.*

### 2. Supplementary Information Results

#### A. Efficiency Analysis

Fig. S2 below outlines the efficiency of the simulation with and without the application of force by comparing the overall runtime to the rate of plotting.

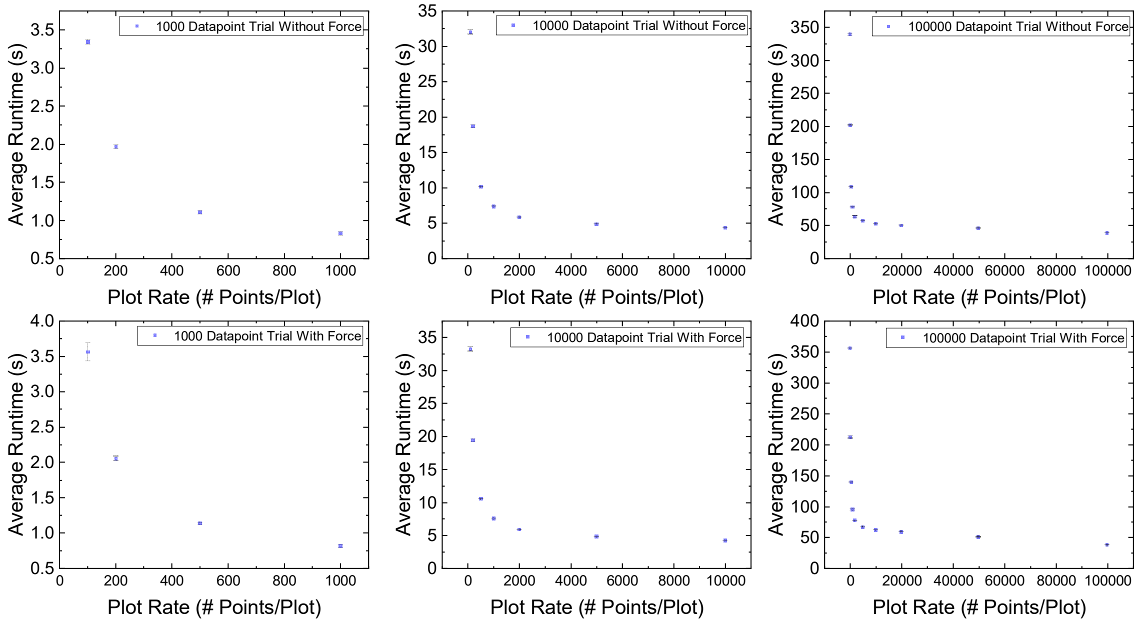


*Fig. S2: Runtime vs. Plot-Rate without Force Ramp (Upper 3) and with Force Ramp (Lower 3)*

The amount of time it takes for data to be generated increases linearly as can be observed from the runtimes that approach the horizontal asymptote in each of the trials above. As the number of plots that must be generated decrease for a specific trial, the runtime can be modeled with an exponentially decaying curve.

Another analysis of the efficiency can be considered using Big-O notation in computer science. This formalism characterizes how efficient an algorithm is through the consideration of the space and time complexities of the algorithm. This simulation is designed using sequential for loops. That is, the for loop which generates the data occurs prior to the for loop which plots the data in the app designer. This means that there are 2 first order complexities in this code since all the implicit computations are computationally constant in time (2). Eq. (i) below provides a general description of the complexity of sequential first order for loops (2).

$$f(n,m) = n * O(1) + m * O(1) \quad \text{(i)}$$

The measure of complexity is done in the limit that the spatial resources approach infinity meaning that some elementary reductions can be made (3):

$$f(n,m) = O(n+m) \quad \text{(ii)}$$

$$f(n,m) = O(\max{(n+m)}) \quad \text{(iii)}$$

The first loop in the sequence generates the data and the second loop plots all the data. This means that n = m yielding Eq. (iv):

$$f(n,n) = O(2n) \quad \text{(iv)}$$

Since the big O notation is a measure of complexity in a limit, pre-factors can be omitted yielding the result (4):

$$f(n) = O(n) \quad \text{(v)}$$

To verify that there is a linear dependence between the data points and runtime, a series of simulations were considered, and runtimes were extrapolated using the inbuilt MATLAB stopwatch. Five repeated trials for each number of data points were conducted and the average runtime was obtained in addition. Verification of linearity can be achieved through the consideration of Pearson's correlation coefficient (5). A linear regression analysis yielded a correlation coefficient of 0.99997. This means that there is a very strong, positive linear relationship between the number of data points in a trial and the runtime in a trial. This indicates that the Big-O analysis was appropriately conducted. A linear dependence in the Big-O formalism corresponds to a decent efficiency.

Overall, there is a point in which the resources necessary to generate the plots continuously greatly exceed the resources necessary to generate the data points. Finding the ideal simulation speed can be done by identifying the data point generation time and finding the point in which the exponential behavior begins dominating. The ideal plot rate will typically exist near this turning point although the total runtime will depend on the specifications of the device used to run the simulation.

#### B. Additional AFS Experimental Comparisons

An additional AFS experiment was conducted to verify the validity of the simulation for a bead size of 2160 nm. In this case, there was a greater asymmetry in the direction of the force application. The generated model only accounts for a force application in the z-direction so it cannot explicitly predict the experimental results with high confidence for an XY rms analysis. Even so, the general trend is captured as is depicted in Fig. S3 below.

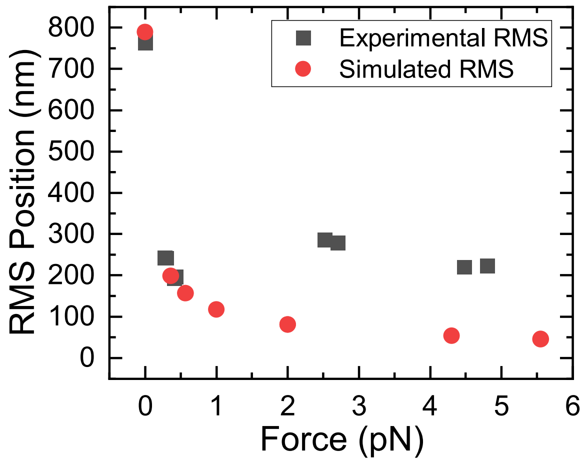


*Fig. S3: Experimental and simulated average XY-RMS position values with application of constant force for 1800 nm DNA strands attached to 2160 nm polystyrene bead in water at room temperature.*

### 3. Supplementary Information Methods

#### A. Experimental Setup of TPM using Acoustic Force Spectroscopy (AFS)

**Chemicals:** 10x Phosphate buffer saline (PBS), Pluronic F-127, casein technical grade, bleach solution, sodium thiosulfate and anti-digoxigenin Fab fragment antibodies (Roche, 11214667001) were purchased from Sigma Aldrich, USA. Bovine serum albumin, fraction V (BSA) was purchased from Goldbio, USA. Streptavidin-coated polystyrene particles with a nominal diameter of 2.16 μm (SVP-20) and 3.11 μm (SVP-30) were purchased from Spherotech Inc, USA.

**Buffers.** All AFS experiments were carried out in working buffer (WB) containing 10 mM PBS at pH 7.4 supplemented with 0.31 mg/ml BSA and casein and 0.19 mg/ml Pluronic F-127, respectively. In addition, two blocking buffers were used to passivate the surface before the experiment. Buffer B1 consists of 10 mM PBS supplemented with 2.5 mg/ml BSA and casein. Buffer B2 consists of 10 mM PBS supplemented with 2.2 mg/ml BSA and casein and 5.6 mg/ml Pluronic F-127 respectively. All buffers were degassed in a vacuum (-90 kPa) for 30 minutes.

**DNA tethers.** Linear double-stranded DNA tethers were synthesized in one step by PCR using the pEC-GFP-CBM3a plasmid (8, 9) as a template and 5' modified primers. The biotin-modified primer (forward primer, 5'-biotin-C6-GGCGATCGCCTGGAAGTA) and digoxigenin modified primer (backward primer, 5'-DIG-NHS- TCCAAAGGTGAAGAACTGTTCACC) were purchased from Integrated DNA Technologies, Inc. USA. The whole plasmid (5.4 kb) was amplified, then purified using the PCR Clean-up kit (IBI Scientific USA) resulting in a linear DNA tether of ~1.8 μm length with one modification on each end of the DNA. Amplification and product purity was verified by gel electrophoresis. For the generation of shorter tethers, a smaller sequence was amplified using the same primer for digoxigenin, and 5'-biotin-C6-GCTTCGAACGCGTATCGATG. With this method, a tether of any size up to the plasmid size can easily be produced.

**Tethered bead preparation for single-molecule force spectroscopy.** Single-molecule experiments were carried out on a G1 AFS instrument with G2 AFS chips provided by LUMICKS B.V. After a 0.5 ml rinse of bleach followed by neutralization with 0.5 ml of 0.5M sodium thiosulfate and wash of 2 ml DI water, the chips were incubated for 20 minutes with anti-digoxigenin fab fragments dissolved in PBS (20 μg/ml), where they non-specifically bind to the AFS glass surface. Next, the surface was passivated with B1 and B2 buffer for 15 minutes each and rinsed with WB. The dig-DNA-biotin tethers were diluted to 6 pM in WB. The bead-DNA-construct was prepared as follows. First, 15μl streptavidin-coated beads and DNA tethers were mixed to yield between 5-15 DNA tethers per and incubated on a rotisserie for 30 minutes. Next, the functionalized beads were washed by spinning the sample down on a table-top centrifuge, removing the supernatant, and resuspending in 100 μl WB twice. After the second removal of supernatant, the DNA-bead construct was resuspended in 30 μl WB.

The DNA-bead construct was flushed through the AFS chip and incubated for 15-30 minutes. Non-bound beads were subsequently washed out with WB at a flow rate of 2 μl/min using a syringe pump (New Era Pump Systems Inc., USA). A small force of ~0.2-0.5 pN was applied to speed up the flushing step. After measuring the rupture forces, the chip was rinsed with 100 µl WB, and the next DNA-bead sample was inserted.

**Bead tracking.** Tracking and analysis of the beads were accomplished using the software package provided by LUMICKS, with slight modifications to allow efficient export of traces and associated tethers statistics as well as force-distance curves to a spreadsheet. The procedure for identifying a single-molecule tether, force calibration, and rupture force determination is described in detail elsewhere (6). The beads were tracked at 20 Hz using a 10x magnification objective. The trajectory of the beads without applied force was monitored for 8-10 minutes to determine the point of surface attachment (anchor point). Next, the force on each bead was calibrated by applying a constant amplitude for 2-4 minutes. Typically, 2-3 different amplitude values were used to build the calibration curve between the applied amplitude and effective force on each bead. Single-molecule tethers were identified by the root-mean-square fluctuation ($RMS$) and symmetrical motion ($Sym$) of the bead around the anchor point during the time frame for anchor point determination. Typical values of single-molecule tethers for $RMS$ and $Sym$ are in the range between 700-1000 nm and 1.0-1.3 respectively for 1800 nm tethers and $RMS$ of 300-450 nm for 500nm tethers. During force calibration, the diffusion coefficient of the bead and the force were used as fit parameters. This diffusion coefficient was compared to the diffusion coefficient determined by the Stokes-Einstein relation and was in the range between 0.8-1.2 for single tethers.

#### Supplementary Information References


1. M. D. Wang, H. Yin, R. Landick, J. Gelles, S. M. Block, Stretching DNA with optical tweezers. *Biophys. J.* **72**, 1335–1346 (1997).
2. S. Bae, *Data structures and algorithms* (2019) https:/doi.org/10.1016/b978-0-7506-0813-8.50017-4.
3. P. Danziger, Big O Notation Basics. 2–7 (2010).
4. S. Gayathri Devi, K. Selvam, S. P. Rajagopalan, An abstract to calculate big o factors of time and space complexity of machine code. *IET Conf. Publ.* **2011**, 844–847 (2011).
5. A. Schneider, G. Hommel, M. Blettner, Lineare regressionsanalyse - Teil 14 der serie zur bewertung wissenschaftlicher publikationen. *Dtsch. Arztebl.* **107**, 776–782 (2010).
6. G. Sitters, *et al.*, Acoustic force spectroscopy. *Nat. Methods* **12**, 47–50 (2014).
7. T. Odijk, Stiff Chains and Filaments under Tension. *Macromolecules* **28**, 7016–7018 (1995).
8. M. Hackl, *et al.*, Acoustic Force Spectroscopy Reveals Subtle Differences in Cellulose Unbinding Behavior of Carbohydrate-Binding Modules. *bioRxiv*, 2021.09.20.461102 (2021).
9. B. Nemmaru, *et al.*, Reduced type-A carbohydrate-binding module interactions to cellulose I leads to improved endocellulase activity. *Biotechnol. Bioeng.* **118**, 1141–1151 (2021).
