## Supplementary material for "Multivariable Graphical User Interface for Simulation of Tethered Particle Motion": MATLAB files: getDragCoefXY.m

```
function dragCoefXY = getDragCoefXY(r, z, sita0)
ratio = r/z; % r = sqrt(x^2 + y^2 + z^2) ~ typical definition 
denom = 1 - (9/16)*ratio + (1/8)*ratio^3 - (45/256)*ratio^4 - (1/16)*ratio^5;
dragCoefXY = sita0 / denom;
```
