## Supplementary material for "Multivariable Graphical User Interface for Simulation of Tethered Particle Motion": MATLAB files: getDragCoefZ.m

```
function dragCoefZ = getDragCoefZ(r, z, sita0)
ratio = r/z; %z >= r and  % r = sqrt(x^2 + y^2 + z^2) ~ typical definition 
denom = 1 - (9/8)*ratio + 0.5*ratio^3 - 0.57*ratio^4 + 0.2*ratio^5 + (7/200)*ratio^11 - (1/25)*ratio^12;
dragCoefZ = sita0 / denom;
```
