## Supplementary material for "Multivariable Graphical User Interface for Simulation of Tethered Particle Motion": MATLAB files: getNextTimestepmod.m

```
function nextTimeStepmod = getNextTimestepmod(sita, Lp, Lo, extx,exty,extz, kbT,Fx,Fy,Fz,err) 
        dia=1.6*1e-9; % Diameter DNA        
        Ko = 16*kbT * Lp * dia^-2; % Youngs modulus
        
        % Determining force gradient finding timestep
        dx = 1/((1 - extx/Lo + Fx/Ko)^3);
        dy = 1/((1 - exty/Lo + Fy/Ko)^3);
        dz = 1/((1 - extz/Lo + Fz/Ko)^3);

        numerx=2/(Lo*dx) + (4/Lo);
        denomx= (4*Lp/kbT) + (2/(Ko*dx)) +(4/Ko);
        dFx=numerx/denomx;
        numery=2/(Lo*dy) + (4/Lo);
        denomy= (4*Lp/kbT) + (2/(Ko*dy)) +(4/Ko);
        dFy=numery/denomy;
        numerz=2/(Lo*dz) + (4/Lo);
        denomz= (4*Lp/kbT) + (2/(Ko*dz)) +(4/Ko);
        dFz=numerz/denomz;

        gradient = sqrt(dFx^2 +dFy^2 +dFz^2);
        nextTimeStepmod = (2*err * sita) / gradient; % err ~ error in displacement, depends on system tolerance

        
```
