## Supplementary material for "Multivariable Graphical User Interface for Simulation of Tethered Particle Motion": MATLAB files: Marko_Siggia_Vectorized.m

```
function force = Marko_Siggia_Vectorized (kbT, Lp, Lo, extx,exty,extz, direction, axis, angleTheta, anglePhi,Rb,applied_force,data)
d=1.6*1e-9; % Diameter DNA
Ko = 16*kbT * Lp * d^-2; % Youngs Modulus

% Format:
% exti ~ magnitude of extension in the i'th direction
% funci ~ defining marko_siggia model with given paramters
% tempFunci ~ inputting variable arguments into funci
% tempForcei ~ solving for force numerically

extx = abs(extx); 
funcx=@(extx,Fe) (kbT/Lp)*(1/4*1/((1- (extx/Lo) +(Fe/Ko))^2)-(1/4)+(extx/Lo)-(Fe/Ko))-Fe;
tempFuncx = @(Fe) funcx(extx, Fe);
tempForcex = fzero(tempFuncx,1e-14);

exty = abs(exty);
funcy=@(exty,Fe) (kbT/Lp)*((1/4)*1/((1- (exty/Lo) +(Fe/Ko))^2)-(1/4)+(exty/Lo)-(Fe/Ko))-Fe;
tempFuncy = @(Fe) funcy(exty, Fe);
tempForcey = fzero(tempFuncy,1e-14);

extz = abs(extz-Rb);
funcz=@(extz,Fe) (kbT/Lp)*((1/4)*1/((1- (extz/Lo)+(Fe/Ko))^2)- (1/4) + (extz/Lo)- (Fe/Ko))-Fe;
tempFuncz = @(Fe) funcz(extz, Fe);
tempForcez = fzero(tempFuncz,1e-14);

tempForce=sqrt(tempForcex^2 +tempForcey^2 +tempForcez^2);

if data == 1 
    if abs(tempForce)>1.1*abs(applied_force) && applied_force ~= 0
        tempForce=1.1*applied_force;
    end
    
    if abs(tempForce)<0.9*abs(applied_force) && applied_force ~= 0
        tempForce=0.9*applied_force;
    end
end

if (direction>0)
    tempForce = -1.0*tempForce;
else
    tempForce = 1.0*tempForce;
end

if (axis == 'x')
    tempForce = tempForce*abs(sin(angleTheta)*cos(anglePhi));
end
if (axis == 'y')
    tempForce = tempForce*abs(sin(angleTheta)*sin(anglePhi));
end
if (axis == 'z')
    tempForce = tempForce*abs(cos(angleTheta));
end

force = tempForce;
```
