## Supplementary material for "Multivariable Graphical User Interface for Simulation of Tethered Particle Motion": MATLAB files: TetherForce2.m

```
function [time,forceOut,xOut,yOut,zOut,extOut,fxout,fyout,fzout,thetaOut,phiOut]=TetherForce2(fLaser,alpha,beta,appliedRange,~,datapoints, Lo, Rb, eta, temperature,Lp,x_mem,y_mem,z_mem,ext_mem,f_mem,t_mem,f_memx,f_memy,f_memz,err,data)
%theta: angle of tether-bead system with respect to z axis
%phi: angle of tether-bead system with respect to xy plane


%alpha: angle of force application with respect to z axis
%beta: angle of the force application with respect to xy plane

%Note: Force only applied in z-direction for simulation current state

n  = datapoints;    %number of data points

%%%%Simulated data%%%%
[Xsim,Ysim,Zsim, FullExtension, AngleSimThetaDegree, AngleSimPhiDegree, DNAForce, timeline,DNAForcex,DNAForcey,DNAForcez] = RandWalkSim(n, fLaser,alpha, beta, appliedRange, Lo, Rb, eta, temperature,Lp,x_mem,y_mem,z_mem,ext_mem,f_mem,t_mem,f_memx,f_memy,f_memz,err,data);  %generate simulated data

% Enclosed Function Outputs
% Xsim ~ Array with x values at each timestep
% Ysim ~ Array with y values at each timestep
% Zsim ~ Array with z values at each timestep
% FullExtension ~ Array with tether extension values at each timestep
% Theta and Phi outputs suppressed but implicitly computed
% DNAForce ~ Total force at each timestep
% timeline ~ Array with time value corresponding to each datapoint
% DNAForcex ~ Array with x-component of force at each timestep
% DNAForcey ~ Array with y-component of force at each timestep
% DNAForcez ~ Array with z-component of force at each timestep

% Enclosed Function Inputs
% n ~ number of datapoints
% fLaser ~ array of applied force values at each datapoint
% alpha ~ array of angle of force with respect to z axis
% beta: array of angle of the force with respect to xy plane.
% appliedRange ~ confidence band of Gaussian distribution
% Lo ~ Contour Length
% Rb ~ Bead Radius
% eta ~ viscosity of thermal bath
% temperature ~ temperature of thermal bath
% Lp ~ tether persistence length

% Re-initialization Inputs: For re-intialization of intial conditions in simulation
% x_mem ~ final x position for fixed number of simulated datapoints
% y_mem ~ final y position for fixed number of simulated datapoints
% z_mem ~ final z position for fixed number of simulated datapoints
% ext_mem ~ final extension for fixed number of simulated datapoints
% f_mem ~ final force value for fixed number of simulated datapoints
% t_mem ~ time for fixed number of simulated datapoints to be generated
% f_memx ~ final x force value for fixed number of simulated datapoints
% f_memy ~ final y force value for fixed number of simulated datapoints
% f_memz ~ final z force value for fixed number of simulated datapoints


% Enclosing Function Output ~ Final Simulation Results for given n
time = timeline; %time series
forceOut=DNAForce; 
fxout=DNAForcex;
fyout=DNAForcey;
fzout=DNAForcez;

xOut=Xsim;
yOut=Ysim;
zOut=Zsim;
extOut=FullExtension;
thetaOut=AngleSimThetaDegree;
phiOut=AngleSimPhiDegree;

function [Xsim, Ysim, Zsim, FullExtension, AngleSimThetaDegree , AngleSimPhiDegree, DNAForce, timeline,DNAForcex,DNAForcey,DNAForcez]=RandWalkSim(n,fLaser,alpha, beta,appliedRange, Lo, Rb, eta, temperature,Lp,x_mem,y_mem,z_mem,ext_mem,f_mem,t_mem,f_memx,f_memy,f_memz,err,data)
%This function simulates a 1D random walk in a harmonic potential

% Energy Unit ~ Prefactor can be modified to consider specific average, ideal,etc. 
kbT = (1.38*10^-23) * (temperature + 273);

% Stokes- Einstein relationship (low Reynolds number) (m^2/s)
D = kbT/(6*pi*eta*Rb); 

% drag coefficient magnitude ~ inverse mobility constant from Faxens law for close proximity for surface.
sita0 = 6*pi*eta*Rb;   

% Initialization of dynamic timestep
deltat = getNextTimestepmod(sita0, Lp, Lo, x_mem,y_mem,z_mem, kbT,f_memx,f_memy,f_memz,err); 

% Declaration of arrays recording simulation data 
Xsim=zeros(1,n);
Ysim=zeros(1,n);
Zsim=zeros(1,n);

DNAForce=zeros(1,n);
DNAForcex=zeros(1,n);
DNAForcey=zeros(1,n);
DNAForcez=zeros(1,n);
AnglePhiSim = zeros(1,n);
AngleThetaSim = zeros(1,n);
AngleThetaSim = AngleThetaSim + pi/2;
AngleSimThetaDegree = zeros(1,n);
AngleSimPhiDegree = zeros(1,n);
timeline = zeros(1,n);

FullExtension = zeros(1,n);

% Initialization of simulation based on initial conditions
Xsim(1,1) = x_mem;
Ysim(1,1) = y_mem;
Zsim(1,1) = z_mem;
DNAForce(1,1) = f_mem;
DNAForcex(1,1) = f_memx;
DNAForcey(1,1) = f_memy;
DNAForcez(1,1) = f_memz;
FullExtension(1,1) = ext_mem;
timeline(1,1) = t_mem;

if Xsim(1,1) ~= 0
    AnglePhiSim(1,1) = atan(Ysim(1,1)/Xsim(1,1));
    AngleSimPhiDegree(1,1) = 180*(AnglePhiSim(1,1)/pi);
     
    AngleThetaSim(1,1) = atan(sqrt(Xsim(1,1)^2 + Ysim(1,1)^2) / Zsim(1,1));
    AngleSimThetaDegree(1,1) = 180*(AngleThetaSim(1,1)/pi);
end

% Declaration of cells to store terms from random number generator
brownianTermsX={};
brownianTermsY={};
brownianTermsZ={};

for i=2:n
    % Initializing current system state using memory terms
    fTemp = fLaser(i,1);
    timeline(i) = timeline(i-1) + deltat;
    extensionX = Xsim(i-1);
    extensionY = Ysim(i-1);
    extensionZ = Zsim(i-1);
    
    anglePhi = AnglePhiSim(i-1);
    angleTheta = AngleThetaSim(i-1);
    
    extension = sqrt(extensionX^2 + extensionY^2 + extensionZ^2);

    FullExtension(i) = extension;
    
    % Determining drag coefficients in each direction
    sitaX = getDragCoefXY(Rb, extensionZ, sita0);
    sitaY = sitaX;
    sitaZ = getDragCoefZ(Rb, extensionZ, sita0);  
 
    % Determining characteristic fluctuation scales (standard deviation)
    ldiff   = sqrt(2*D*deltat);
    
    % Generating randon fluctuation terms to account for brownian motion
    bTX = normrnd(0,ldiff,1,1);
    brownianTermsX{i}=bTX;
    bTY = normrnd(0,ldiff,1,1);
    brownianTermsY{i}=bTY;
    bTZ = normrnd(0,ldiff,1,1);
    brownianTermsZ{i}=bTZ;
    
    % Decalration of tether properties (DNA)
    d=1.6*1e-9; % Diameter DNA
    Ko = 16*kbT * Lp * d^-2; % Youngs Modulus

    if fTemp == 0 % Use finite steps, equillibrium assumptions
       
        % Determining force in each direction
        DNAForceX = Marko_Siggia_Vectorized(kbT, Lp, Lo, extensionX,extensionY,extensionZ, extensionX,'x', angleTheta, anglePhi,Rb,fTemp,data);
        DNAForceY = Marko_Siggia_Vectorized(kbT, Lp, Lo, extensionX,extensionY,extensionZ, extensionY,'y', angleTheta, anglePhi,Rb,fTemp,data);
        DNAForceZ = Marko_Siggia_Vectorized(kbT, Lp, Lo, extensionX,extensionY,extensionZ, extensionZ,'z', angleTheta, anglePhi,Rb,fTemp,data);
        
        % Modyifying based on force application orientation
        % Note: fixed to z direction only for this simulation

        forceX = DNAForceX + fTemp*sin(alpha(i,1))*cos(beta(i,1)); %  N Scaling
        forceY = DNAForceY + fTemp*sin(alpha(i,1))*sin(beta(i,1)); 
        forceZ = DNAForceZ + fTemp*cos(alpha(i,1));

        DNAForcex(1,i) = forceX; % Converting to N scaling
        DNAForcey(1,i) = forceY;    
        DNAForcez(1,i) = forceZ;

        DNAForce(1,i) = sqrt(DNAForceX^2 + DNAForceY^2 + DNAForceZ^2);
        
        % Updating position elements

        deltaX= (forceX*deltat)/sitaX + brownianTermsX{i};
        Xsim(i)=Xsim(i-1)+deltaX;%new element value  
    
        deltaY= (forceY*deltat)/sitaY + brownianTermsY{i};    
        Ysim(i)=Ysim(i-1)+deltaY; %new element value 

        deltaZ= (forceZ*deltat)/sitaZ + brownianTermsZ{i};
    
        if (Zsim(i-1) + deltaZ < Rb)
            Zsim(i)=Rb; %new element value   
        else
            Zsim(i)=Zsim(i-1)+deltaZ;
        end
    end
    % Initiating probabilistic planar, artificial equillibrium using two
    % models. This model is specifically for trend analysis and has a clear
    % transition point between probabilistic and maximization approach.
    % Further details can be found in manuscript.
    
    %data=0 implies probabilistic with transition to maximization

    if fTemp ~= 0 && data==0 

        DNAForceX = Marko_Siggia_Vectorized(kbT, Lp, Lo, extensionX,extensionY,extensionZ, extensionX,'x', angleTheta, anglePhi,Rb,fTemp,data);
        DNAForceY = Marko_Siggia_Vectorized(kbT, Lp, Lo, extensionX,extensionY,extensionZ, extensionY,'y', angleTheta, anglePhi,Rb,fTemp,data);
        DNAForceZ = Marko_Siggia_Vectorized(kbT, Lp, Lo, extensionX,extensionY,extensionZ, extensionZ,'z', angleTheta, anglePhi,Rb,fTemp,data);

        forceX = DNAForceX + fTemp*sin(alpha(i,1))*cos(beta(i,1)); %  N Scaling
        forceY = DNAForceY + fTemp*sin(alpha(i,1))*sin(beta(i,1)); 
        forceZ = DNAForceZ + fTemp*cos(alpha(i,1));

        DNAForcex(1,i) = forceX; % Converting to N scaling
        DNAForcey(1,i) = forceY;    
        DNAForcez(1,i) = forceZ;

        DNAForce(1,i) = sqrt(DNAForceX^2 + DNAForceY^2 + DNAForceZ^2);

       deltaX= (forceX*deltat)/sitaX + brownianTermsX{i};
       Xsim(i)=Xsim(i-1)+deltaX;%new element value  
         
       deltaY= (forceY*deltat)/sitaY + brownianTermsY{i};    
       Ysim(i)=Ysim(i-1)+deltaY; %new element value
            
       Fez = abs(fTemp*cos(alpha(i,1))); % System forced into equillibrium ~ Applied Force dominates the behavior
       funcz=@(extensionZ, Fez) kbT/Lp*(1/4*1/(1-extensionZ/Lo+Fez/Ko)^2-1/4+extensionZ/Lo-Fez/Ko)-Fez;
       tempFuncz = @(extensionZ) funcz(extensionZ, Fez);
       z_eq=fzero(tempFuncz,Lo);
            
       deltaZ = z_eq+Rb +normrnd(0,(Lo-z_eq)/3,1,1); % Generates distribution about new equillibrium state
            
       if deltaZ<Rb
          Zsim(i)=Rb;
       else
          Zsim(i)=deltaZ;
       end
    
       % Initiating planar maximization + resuming normal artificial equillibrium   
    
       if (Xsim(i)^2 +Ysim(i)^2 +Zsim(i)^2)^0.5 > (Lo+Rb)
           range = (Lo+Rb)-abs(deltaZ);
           optimal_ratio=abs(forceX/forceY); % Fx/Fy ~ x/y
           x_sign=forceX/abs(forceX);
           y_sign=forceY/abs(forceY);
           y_bound=range;
           deltaY = y_sign*(y_bound)/(optimal_ratio +1);
           Ysim(i)=(deltaY)+ normrnd(0,(Lo-z_eq)/3,1,1);% Maintaining assumption of isotropy when system forced into equillibrium
    
           deltaX=x_sign*(abs(range-deltaY));
           Xsim(i)=(deltaX)+normrnd(0,(Lo-z_eq)/3,1,1);% Maintaining assumption of isotropy when system forced into equillibrium

        end
    end
    
    %data=1 implies maximization throughout
    if data==1 && fTemp~=0 % For data strictly
        DNAForceX = Marko_Siggia_Vectorized(kbT, Lp, Lo, extensionX,extensionY,extensionZ, extensionX,'x', angleTheta, anglePhi,Rb,fTemp,data);
        DNAForceY = Marko_Siggia_Vectorized(kbT, Lp, Lo, extensionX,extensionY,extensionZ, extensionY,'y', angleTheta, anglePhi,Rb,fTemp,data);
        DNAForceZ = Marko_Siggia_Vectorized(kbT, Lp, Lo, extensionX,extensionY,extensionZ, extensionZ,'z', angleTheta, anglePhi,Rb,fTemp,data);

        % alpha,beta ~ orientation: fixed at alpha=beta=0 for this
        % simulation
        forceX = DNAForceX + fTemp*sin(alpha(i,1))*cos(beta(i,1)); %  N Scaling
        forceY = DNAForceY + fTemp*sin(alpha(i,1))*sin(beta(i,1)); 
        forceZ = DNAForceZ + fTemp*cos(alpha(i,1));

        DNAForcex(1,i) = forceX; 
        DNAForcey(1,i) = forceY;    
        DNAForcez(1,i) = forceZ;

        DNAForce(1,i) = sqrt(DNAForceX^2 + DNAForceY^2 + DNAForceZ^2);

       % Arbitrary initialization of position values to start simulation
       Xsim(i)=1e-9;  
       Ysim(i)=1e-9; 
       
       % System forced into equillibrium ~ Applied Force dominates the behavior
       Fez = abs(fTemp*cos(alpha(i,1))); 
       funcz=@(extensionZ, Fez) kbT/Lp*(1/4*1/(1-extensionZ/Lo+Fez/Ko)^2-1/4+extensionZ/Lo-Fez/Ko)-Fez;
       tempFuncz = @(extensionZ) funcz(extensionZ, Fez);
       z_eq=fzero(tempFuncz,Lo);
            
       deltaZ = z_eq+Rb +normrnd(0,(Lo-z_eq)/3,1,1); % Generates distribution about new equillibrium state
            
       if deltaZ<0
          Zsim(i)=0;
       else
          Zsim(i)=deltaZ;
       end

         if  i>2 % not initial state
           
           if deltaZ>Lo
               deltaZ=Lo;
           end
          % Defining reasonable span of values
           range = abs(Lo-abs(deltaZ));

           % Maintaining assumption of isotropy when system forced into equillibrium
           % accounting for extensibility within 3 degrees of freedom

           Ysim(i)=range/2 + normrnd(0,(Lo-z_eq)/3,1,1); 
           Xsim(i)=range/2 + normrnd(0,(Lo-z_eq)/3,1,1);

        end
        
    end
    
    % Updating theta,phi positions based on positions in cartesian system
    AnglePhiSim(i) = atan(Ysim(i)/Xsim(i));
    AngleSimPhiDegree(i) = 180*(AnglePhiSim(i)/pi);
     
    AngleThetaSim(i) = atan(sqrt(Xsim(i)^2 + Ysim(i)^2) / Zsim(i));
    AngleSimThetaDegree(i) = 180*(AngleThetaSim(i)/pi);
     

    if (sitaX < sitaZ)
        sita = sitaX;
    else 
        sita = sitaZ;
    end

    deltat = getNextTimestepmod(sita, Lp, Lo, extensionX,extensionY,abs(extensionZ-Rb), kbT,DNAForceX,DNAForceY,DNAForceZ,err);
    deltat=abs(deltat);

end
```
