## Supplementary figures and images for "Multivariable Graphical User Interface for Simulation of Tethered Particle Motion"

### TPM Updated Sketch.png

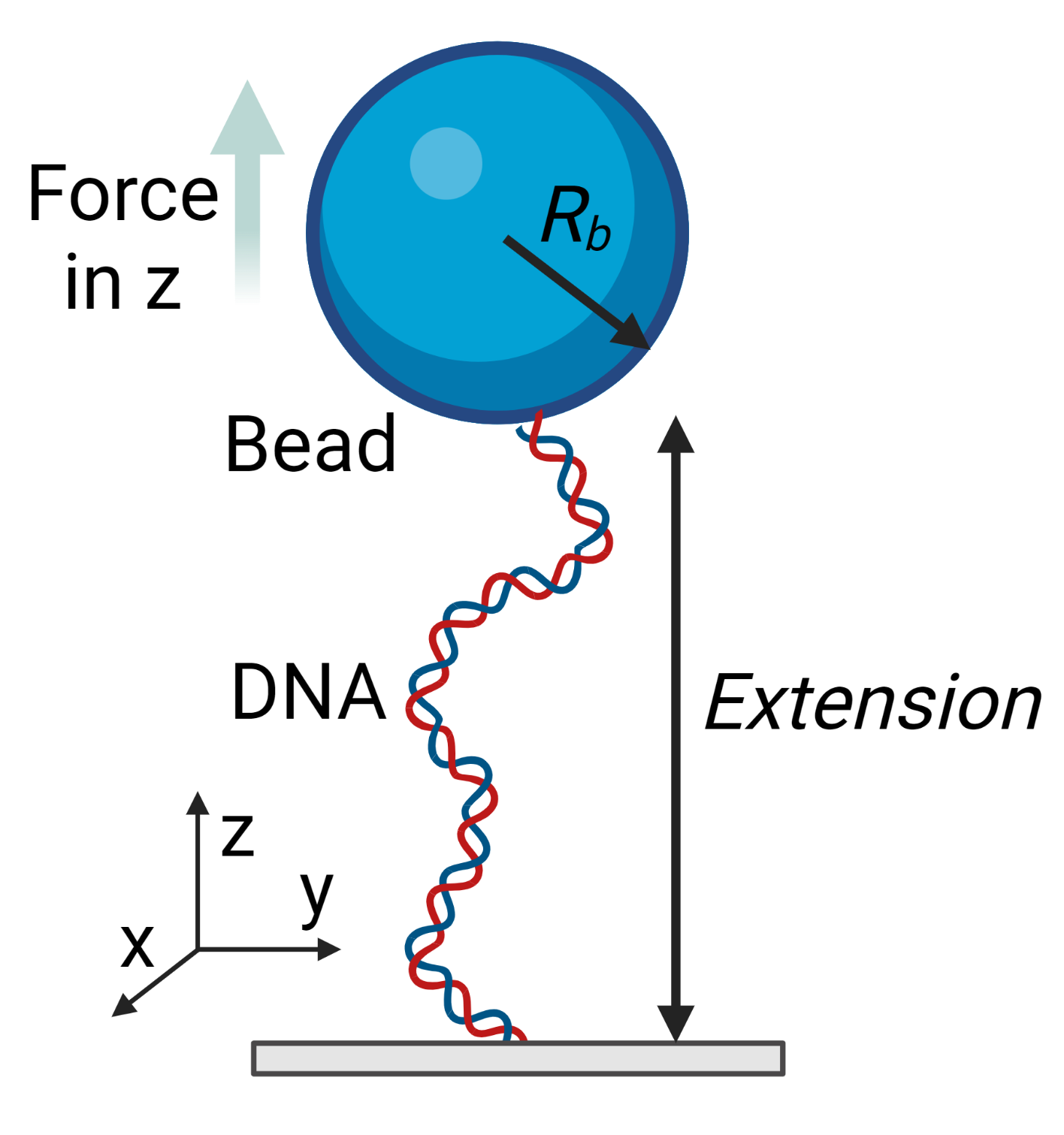
